# Behavioural Profiling of Therapeutic Antibodies via Non-equilibrium Interfacial Wave Dynamics

**DOI:** 10.1101/2025.05.09.653084

**Authors:** Alec Nelson Thomas, Alexander Nicholas St John, Maureen Crames, Piergiorgio Caramazza, Alireza Meghdadi, Joe Bailey, Rodolfo Hermans, Sejeong Lee, Elizabeth Lam, Vivek Krishnan, Alexandra Amoruso, Sam Aoudjane, Michael S. Marlow, Carolina Zirn, Pratik Kotkar, Akshay Mishra, Anshika Srivastava, Shamit Shrivastava

## Abstract

Therapeutic antibody performance depends not only on sequence and structure, but also on how the molecule responds to physico-chemical perturbations encountered in its environment. Whereas sequence and static structure can be inferred under controlled conditions, behaviour is a conditional, multidimensional response to environment, most directly characterised by applying defined perturbations and quantifying the resulting dynamics. Conventional early-stage developability methods capture isolated dimensions of this response under near-equilibrium conditions, deferring integrated behavioural assessment to late-stage characterisation, when the cost of correction is highest.

We introduce Variations in Interfacial Behaviour under Excitation (VIBE), an interfacial wave method implemented via Liquid State Intelligence (LSI), a sensing architecture that transduces molecular perturbations at the air–liquid interface into high-dimensional wave patterns. A colloidal liquid substrate operated near a thermodynamic transition couples small molecular perturbations to large dynamical responses, integrating structural flexibility, charge distribution, and surface hydrophobicity into a single behavioural readout from microgram-scale samples.

Applied to antibodies previously characterised by industrial benchmarks, the primary behavioural descriptor, VIBE1, functioned as a high-precision triage tool, flagging candidates carrying multiple biophysical liabilities. In a clinical-stage cohort, the proportion of high-VIBE1 antibodies declined progressively from early trials through approval, and high-VIBE1 candidates showed an elevated clinical failure rate. Concordance analysis against established methods confirmed that VIBE1 captures a composite signal spanning hydrophobicity, polyreactivity, self-interaction, and thermal stability rather than recapitulating any single conventional readout. These findings establish interfacial wave sensing as a low-material modality for early-stage developability assessment, repositioning molecular behaviour from late-stage validation to discovery-phase characterisation.

## Introduction

Antibody performance depends not only on sequence and structure, but also on how the molecule responds to physico-chemical perturbations encountered in its environment. When antibodies encounter dynamic interfaces such as the air–water boundary, adsorption, deformation, and intermolecular interactions can generate behaviours that are not evident under near-equilibrium bulk conditions. Whereas sequence and static structure can often be inferred under controlled conditions, behaviour is a conditional, multidimensional response to the molecule’s environment and is most directly characterised by applying defined, context-relevant perturbations and quantifying the resulting dynamics [31]. This asymmetry, together with constraints of throughput and material availability, has tended to defer behavioural investigation to later development stages. Specifically, in therapeutic antibody development, sequence is generated at discovery and structure may be solved during lead optimisation; isolated developability flags can be obtained early [2], but integrated behavioural assessment under the dynamic stresses encountered in manufacturing and formulation typically requires sufficient material to defer it to late-stage characterisation, when the cost of correction is highest [1, 36, 37].

The standard repertoire of early-stage developability methods—spanning hydrophobicity (e.g. Hydrophobic Interaction Chromatography, HIC), polyreactivity (e.g. Baculovirus Particle ELISA, BVP), self-interaction (e.g. Self-Interaction Nanoparticle Spectroscopy, SINS), and thermal stability (e.g. Differential Scanning Fluorimetry, DSF)—each captures an isolated dimension of molecular behaviour under controlled, near thermodynamic transition conditions [3, 4]. Individual laboratories typically deploy a subset of these methods; no single panel is universal.

While individually informative, these methods provide limited insight into the complex, time-dependent biophysics that emerge when antibodies encounter real-world stresses such as air–water interfaces, shear, freeze–thaw cycles, and excipient-induced crowding [8, 26, 27, 28, 3]. The critical metric for developability may not be any single property, but rather the integrated molecular behaviour under stress.

As a consequence, the most informative behavioural data – how a molecule responds to the stresses of manufacturing, formulation, and storage – becomes available only after substantial investment in expression, purification, and scale-up. In early antibody discovery, candidates are first selected from screening libraries at hit identification and subsequently refined during lead optimisation. Decisions at these stages, where the candidate pool is largest and the cost of correction lowest, are therefore made with limited behavioural visibility [2]. This temporal mismatch between when behavioural information is needed and when it becomes measurable defines the central challenge addressed here.

Interfacial methods such as the Langmuir trough and pendant-drop tensiometry can probe molecular behaviour at surfaces, but established techniques occupy different corners of a three-way trade-off between data quality, material efficiency, and measurement speed. Langmuir-trough isotherms provide rich thermodynamic information but require large surface areas, milligram quantities, and quasi-static compression over hours [5, 6, 7]; pendant-drop and bubble methods are faster and more material-efficient but resolve only a small number of interfacial observables per measurement. For screening therapeutic candidates at hit identification, where material is limiting, turnaround must be rapid, and a multi-dimensional behavioural readout is desired, established methods have not delivered this combination [8].

We have developed an alternative approach, termed Liquid State Intelligence (LSI) [11, 10, 9, 15], that replaces observations of quasi-static reversible phenomena with repeated non-equilibrium surface excitations. The substrate is a colloidal liquid maintained near its critical micellisation concentration, a regime in which it sits close to a thermodynamic phase transition: deposited sample molecules with surface activity partition strongly to the air–liquid boundary (local concentration enrichment), and small perturbations couple to large dynamical responses (stress-probe sensitivity) [13, 33, 14]. Sample droplets impose mechanical deformation and chemical gradients in parallel [41, 40]; the resulting interfacial wave response, combining capillary and gravity modes, is recorded via an ellipsometric optical setup [6, 17, 18] at approximately 10 *µ*s resolution.

Each measurement begins with a droplet. During droplet formation a new interface is created that then coalesces with the substrate, mixing the droplet’s buffer with the substrate environment (driving abrupt changes in ionic and excipient conditions for the protein) while pinching of the droplet imposes elongational stress on the molecules; subsequent mixing and convection update the global state of the substrate before the cycle repeats. This sequence subjects sample molecules to a complex trajectory through thermodynamic state space, eliciting behavioural responses dependent on their physicochemical properties. Following Eigen’s chemical relaxation approach [19], the system is perturbed from equilibrium and its return is recorded; the resulting relaxation yields kinetic information and can resolve rich multi-step mechanisms that equilibrium measurements alone do not explore. The droplet’s impact transfers momentum to the substrate surface, encoding the properties of the droplet interface into waves propagating across it. The dispersion, nonlinearity, and dissipation of these waves during propagation stretch the hitherto compressed dynamical information into signals resolvable by simpler instrumentation than would otherwise be needed, a principle inspired by physical reservoir computing architectures [34, 35]. Over approximately 100 droplets (~400 s), the measurement follows a five-step actuation cycle (Figure 1A): droplet growth with protein adsorption to the expanding interface, coalescence with the sensing liquid, wave propagation through the liquid film, waveform capture, and repetition — tracking behavioural evolution as sample accumulates at the point of deposition. By applying a hierarchical moment analysis (detailed in Materials and Methods), we compress the resulting high-bandwidth temporal data into per-sample numerical descriptors (one VIBE feature per antibody). The top-ranked representative, VIBE1, quantifies the consistency of the interfacial waveform shape across successive droplet depositions during early titration. The behavioural fingerprint generated by this approach is not designed to isolate any single biophysical mechanism, but to capture composite responses whose predictive value is established empirically by correlation with developmental outcomes. Here, we benchmark VIBE1 on therapeutic antibody developability screening. In an industrial cohort, VIBE1 achieves 100% precision for molecules with multiple biophysical flags and across 235 clinical-stage antibodies, it achieves 85% precision in identifying candidates that did not achieve regulatory approval, using less than 10 *µ*g per measurement.

**Figure 1.**
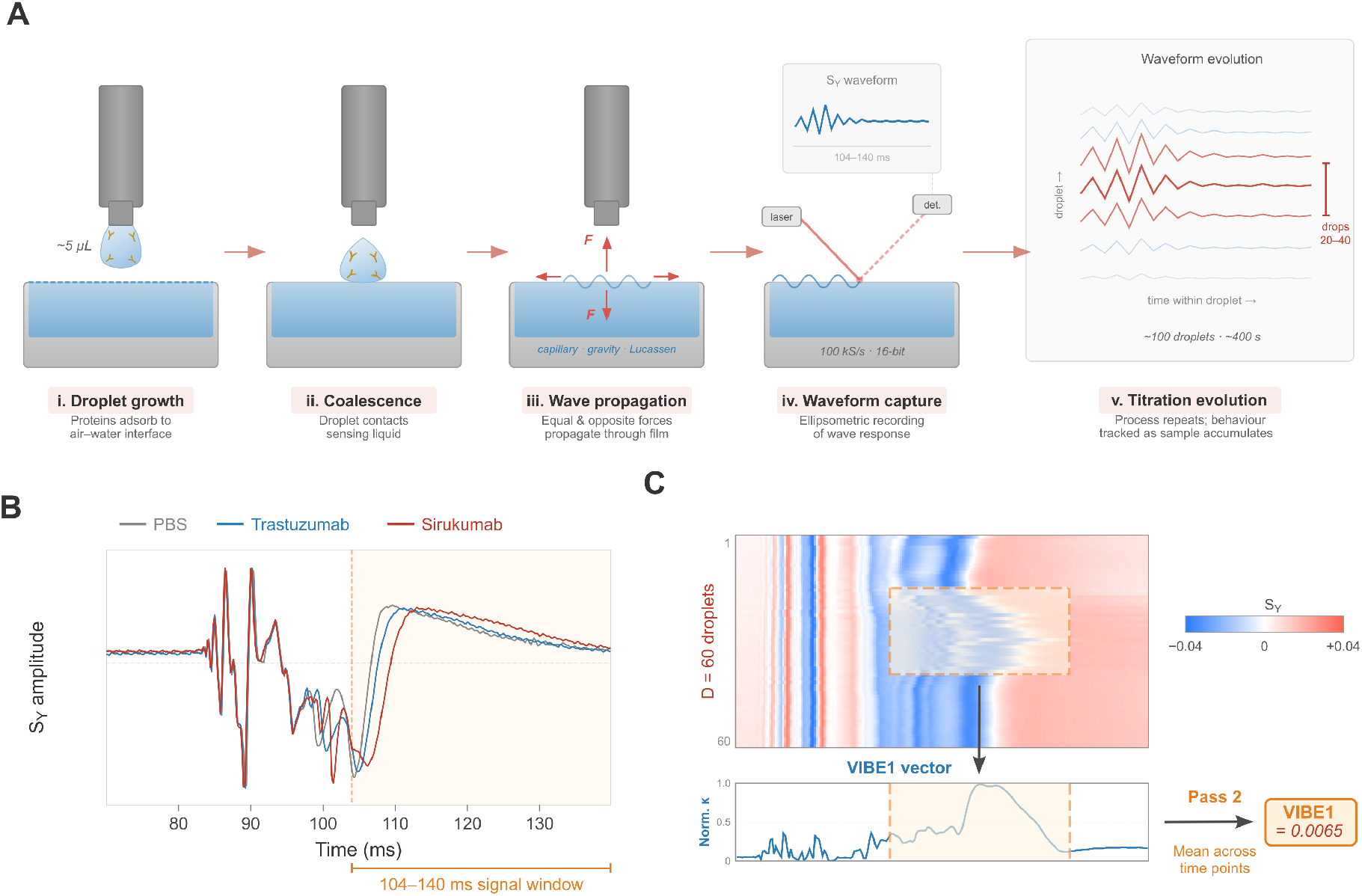
The VIBE method: interfacial wave sensing via the LSI platform. (*A*) Five-step measurement sequence: droplet growth, coalescence, wave propagation, waveform capture, and titration evolution. (*B*) Representative waveforms from the *S*_*Y*_ ellipsometric channel showing sample-specific interfacial responses in the 104–140 ms wave packet (PBS, trastuzumab, sirukumab). (*C*) Two-pass hierarchical moment analysis: the waveform matrix (droplets × time points) is compressed by column-wise kurtosis with *n*-th root standardisation, followed by averaging across time points, yielding the VIBE1 scalar descriptor.

## Results

Two complementary antibody cohorts were used to develop and evaluate VIBE1. Dataset 1 comprises 71 clinical antibodies provided by Boehringer Ingelheim, for which five industry-standard developability methods had been performed in-house, providing direct biophysical-flag labels. Dataset 2 comprises 235 clinical-stage antibodies sourced from Thera-SAbDab [20], with known approval status and phase-progression history but no in-house biophysical-method results. Together these cohorts provide independent dimensions of validation: Dataset 1 anchors VIBE1 to specific biophysical liabilities, while Dataset 2 tests whether the resulting risk signal carries through to clinical outcome.

### Observation-driven descriptor design and VIBE1 selection

We observed qualitative waveform differences between control antibodies sirukumab and trastuzumab (Figure 1B) concentrated around droplets 20–40, corresponding to early-phase interfacial accumulation. This motivated a general descriptor framework, VIBE, based on a two-pass hierarchical moment analysis (Eqs. S1–S4).

To identify the most discriminating descriptor, we performed a label-blind supervised feature selection across 320 candidate features spanning four optical channels, five signal windows, and 16 moment-pair combinations. Features were ranked by ANOVA *F*-statistic on biophysical-flag class assignments, a criterion that does not see clinical outcome, and the top-ranked descriptor was selected for downstream analysis. The procedure was designed to be robust to specific parameter choices: high-*F* features formed a tightly correlated cluster (SI Appendix, Figure S1) indicating that a broad family of features captures the same physical signal rather than a single fragile descriptor, and the same descriptor topped the ranking on both Dataset 1 and Dataset 2 (next paragraph).

The top-ranked feature, designated VIBE1, is computed from the *S*_*Y*_ channel over the 104–140 ms signal window (Figure 1C): the 4th central moment across droplets (first pass), root-standardised, then averaged across time points (second pass). VIBE1 quantifies how consistently the interfacial waveform evolves across successive depositions during early titration; high values indicate strong drop-to-drop fluctuation. This feature was identified as the top-ranked by *F*-statistic in both Dataset 1 and Dataset 2.

### VIBE1 identifies composite biophysical risk with high precision

To benchmark VIBE1 as a behavioural readout, we evaluated a panel of 71 clinical antibodies (Dataset 1) for which five industry-standard developability methods had been performed: purity after Protein A capture, measured by SEC (% purity), Fab melting temperature (Fab *T*_*m*_), analytical hydrophobic interaction chromatography (aHIC), self-interaction by SINS, and non-specific binding (NSB) [3, 21, 22]. Antibodies exceeding the thresholds defined in Table 1 on two or more methods were categorised as high-risk; these thresholds follow industry conventions reported by [3, 21, 22], applied at Boehringer Ingelheim independently of the VIBE1 measurement. Of the 71 antibodies, 18 exhibited two or more conventional flags.

**Table 1.** Biophysical risk thresholds for Dataset 1, following industry conventions [3, 21, 22] and applied at Boehringer Ingelheim independently of the VIBE1 measurement. Antibodies classified as “High Risk” on two or more methods were designated as composite high-risk candidates.

| | Purity % | Fab $T_m$ | aHIC | SINS | NSB |
| --- | --- | --- | --- | --- | --- |
| Optimal | $\geq 95$ | $\geq 78$ | $< 6.0$ | $\geq 1.68$ | $\leq 1.25$ |
| Normal | $\geq 90$ | $\geq 72$ | $\leq 10$ | $\geq 1.63$ | $< 1.8$ |
| High Risk | $< 90$ | $< 72$ | $> 10.0$ | $< 1.63$ | $\geq 1.8$ |

**Abbreviations:** % SEC, purity after Protein A capture by SEC (% purity); Fab $T_m$ , Fab melting temperature ( $^{\circ}\text{C}$ ) by DSF; aHIC, analytical hydrophobic interaction chromatography retention; SINS, self-interaction nanoparticle spectroscopy; NSB, non-specific binding score.

To evaluate VIBE1 as a biophysical risk classifier, a VIBE1 flagging threshold of *τ*_1_ = 0.036 which had previously been identified by fitting a Gaussian mixture model (GMM) to Dataset 2 (see Materials and Methods) was applied to Dataset 1 after min–max rescaling against sirukumab and trastuzumab internal controls. A *de novo* GMM fitted directly to Dataset 1 produced an almost identical threshold (*τ*_1_ = 0.037) which resulted in no difference in classification performance, confirming cross-dataset consistency. Sirukumab, the high developability risk control consistently shows higher VIBE1 score than the low risk control, trastuzumab. Therefore, antibodies above the threshold were classed as high-risk, and antibodies below the threshold were classed as low-risk.

Using the GMM threshold, VIBE1 flagged 6 antibodies as high-risk; all 6 were confirmed multi-flag molecules, yielding a precision of 1.00 (95% CI: 0.61–1.00; Figure 2A,B), with 53 true negatives, zero false positives, 12 false negatives, and recall of 0.33 (95% CI: 0.16–0.56).

**Figure 2.**
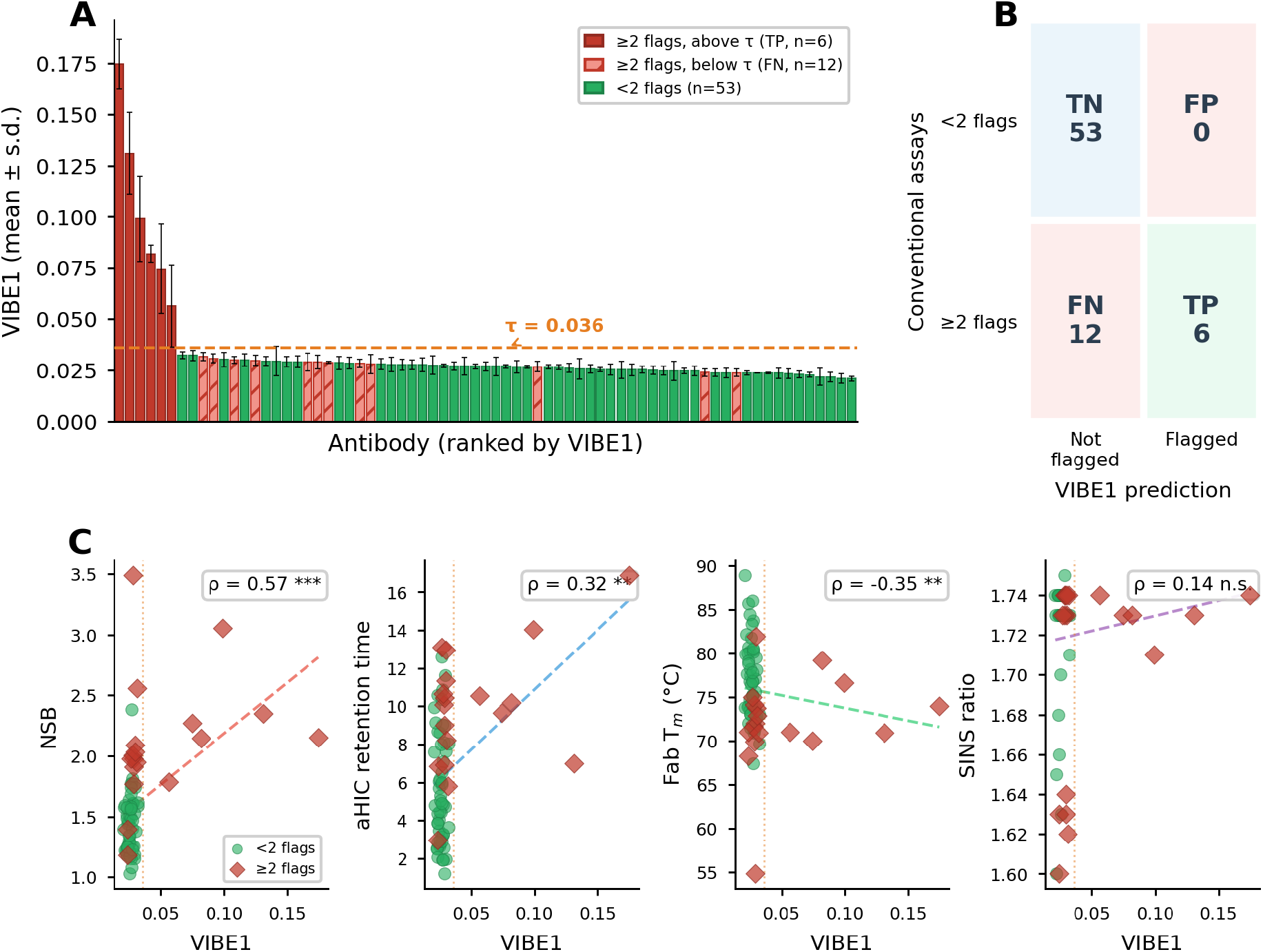
VIBE1 identifies multi-flag antibodies with 100% precision and correlates with multiple biophysical methods. (*A*) VIBE1 values for 71 antibodies in the Boehringer Ingelheim cohort, ranked by magnitude. Red bars: antibodies with ≥ 2 conventional flags above threshold (true positives); hatched bars: ≥2 flags below threshold (false negatives); green bars: *<*2 flags. Dashed line: classification threshold (*τ*_1_ = 0.036), transferred from Dataset 2 (see Materials and Methods). Error bars: ±s.d. across replicate measurements. (*B*) Confusion matrix. TN = true negative; FP = false positive; FN = false negative; TP = true positive. (*C*) Spearman rank correlations between VIBE1 and four conventional biophysical methods. Points coloured by multi-flag status (green: *<*2 flags; red diamonds: ≥2 flags). Dotted vertical line indicates GMM threshold. Dashed lines: linear trend for visualisation. Significance: \*\*\**p <* 0.001; \*\**p <* 0.01; n.s. = not significant.

### VIBE1 captures composite behavioural information

Correlation analysis between VIBE1 and the four conventional methods revealed patterns consistent with the LSI measurement being sensitive to underlying biophysical properties and integrating information across multiple dimensions (Figure 2C). VIBE1 showed a moderate positive correlation with NSB (*ρ* = 0.57, *p <* 0.001), a weaker positive correlation with aHIC retention time (*ρ* = 0.32, *p* = 0.006), and a negative correlation with Fab *T*_*m*_ (*ρ* = −0.35, *p* = 0.003). No significant correlation was observed with SINS ratio (*ρ* = 0.14, *p* = 0.23).

These varied correlations suggest that VIBE1 captures a composite behavioural signal sensitive to multiple biophysical properties rather than serving as a proxy for any single conventional method. The strongest association—with NSB—is consistent with the interfacial nature of the measurement, where surface-active molecules that exhibit non-specific binding also perturb the liquid substrate more strongly. The positive correlation with hydrophobicity (aHIC) and negative correlation with thermal stability (*T*_*m*_) are consistent with VIBE1 being preferentially responsive to molecules carrying multiple concurrent liabilities (hydrophobic, electrostatic, and conformationally unstable), although the present data demonstrate enrichment of multi-liability molecules among VIBE1-flagged candidates rather than a multiplicatively synergistic response. This behaviour is consistent with VIBE1 resolving liabilities that may be attenuated or obscured under the static conditions of conventional screening panels.

### Clinical pipeline validation: VIBE1 enriches for failure risk

If VIBE1 captures biophysical liabilities relevant to clinical attrition, then it should enrich for the molecules that ultimately fail in development. We tested this prediction on a larger independent cohort: 235 clinical-stage antibodies (Dataset 2) with known approval status and phase-progression history, sourced from Thera-SAbDab [20]. Experiments were performed on the LSI using the same hardware configuration as Dataset 1. Due to differences in supplier and buffer (PBS rather than histidine), the absolute VIBE1 scores for control antibodies differed from Dataset 1, consistent with VIBE1 sensitivity to all components of the droplet.

Because Dataset 2 was acquired first and spans a larger, more diverse clinical cohort, the classification threshold was defined *de novo* on this dataset. A two-component GMM fitted to replicate-averaged VIBE1 values identified a bimodal structure and yielded *τ*_2_ = 0.041, obtained without fitting to the target variable (see Materials and Methods; SI Appendix).

The baseline non-approval rate was 72.3% (65/235 approved; 95% CI: 66.3–77.7%). VIBE1 flagged 41/235 antibodies (17.4%; 95% CI: 13.1–22.8%) as high risk. Among flagged antibodies, 35/41 were not approved (85.4%; 95% CI: 71.6–93.1%), corresponding to an absolute increase in non-approval of 13.1 percentage points relative to baseline (95% CI: −6.1–26.8). Recall for non-approval (flagged among not approved) was 20.6% (95% CI: 15.2–27.3%), indicating an operating point that prioritises precision over recall, consistent with Dataset 1.

We next asked whether VIBE1 flags would track development progression: would flagged antibodies become rarer at later stages? The fraction of high-VIBE1 antibodies declined monotonically from Phase I/II as candidates progressed towards regulatory approval (Figure 3A): Phase I, 20.7%; Phase II, 22.0%; Phase III, 14.3%; Approved, 9.2%. Furthermore, among antibodies still actively in clinical trials or approved at the time of analysis, 86.8% exhibited low VIBE1 scores. This progressive filtering where high-VIBE1 molecules are systematically underrepresented at later stages is consistent with behavioural outliers being preferentially discontinued at each stage, or never advanced as frequently. Both interpretations support the inference that high VIBE1 scores associate with characteristics that impede successful development.

**Figure 3.**
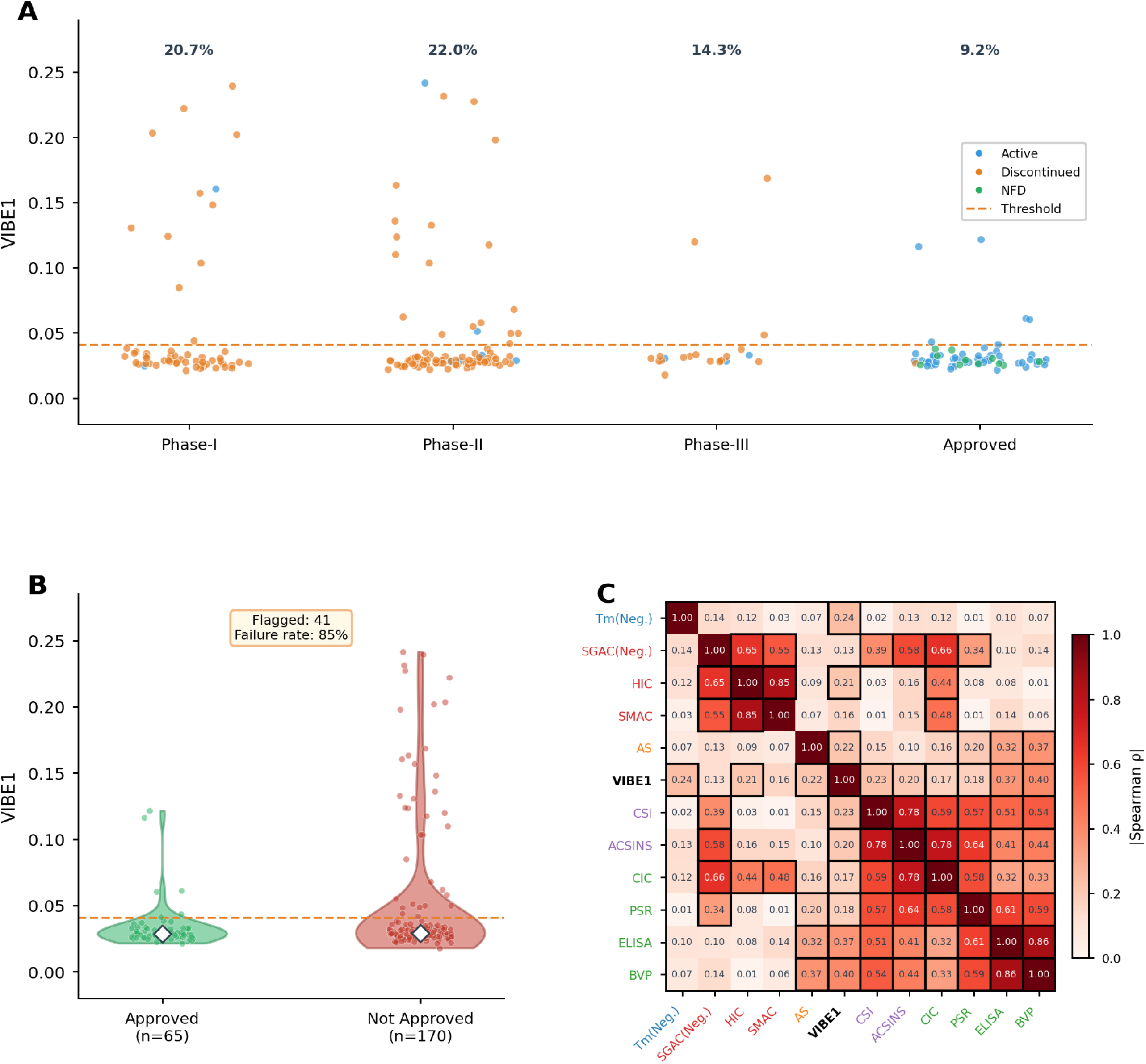
Clinical pipeline validation of VIBE1. (*A*) VIBE1 scores by highest clinical stage reached (*n* = 235). Dashed line: risk threshold (*τ* = 0.041). Percentages: fraction above threshold per stage. Points coloured by current development status (blue: active; orange: discontinued; green: no further development). (*B*) Distribution of VIBE1 scores for approved (*n* = 65) versus non-approved (*n* = 170) antibodies. Diamonds indicate medians. (*C*) Spearman rank correlation heatmap (absolute values) comparing VIBE1 scores from 135 IgG1-format antibodies (82 native IgG1 plus 53 re-expressed) from Dataset 2 with 11 conventional methods from Jain et al. [21] (*n* = 135). Tick labels coloured by property class: hydrophobicity (red), polyreactivity (green), self-interaction (purple), thermal stability (blue), aggregation (orange), VIBE (black). Black borders outline statistically significant pairs (*α <* 0.05). The modest pairwise correlations indicate that VIBE1 captures information independent of any single existing method; despite this independence, VIBE1 performs comparably to several widely used methods in discriminating clinical outcome (see main text).

Finally, we asked which biophysical properties VIBE1 most resembles when compared head-to-head against an established panel. Among 135 IgG1-format antibodies (82 native IgG1 plus 53 re-expressed) measured for like-for-like comparison with the eleven-method panel of Jain et al. [21], VIBE1 showed moderate positive correlation with two polyreactivity methods BVP (*ρ* = 0.40), ELISA (*ρ* = 0.37), with weaker associations (*ρ* = 0.13 − 0.24) across hydrophobicity, thermal stability, self-interaction, and aggregation methods (Figure 3C). In hierarchical clustering, VIBE1 positioned between the hydrophobicity and polyreactivity clusters and showed more statistically significant correlations than any other method.

### Structural correlates of the VIBE1 signal

To explore whether structure-derived features encode information about the VIBE1 signal, we trained a Random Forest classifier on DeepSP [23] spatial aggregation propensity (SAP) and spatial charge map (SCM) descriptors across the full 235-antibody cohort. The model predicted VIBE1 classification with an ROC-AUC of 0.766 (95% CI: 0.665–0.851; SI Appendix, Figure S3). SHAP analysis [38] identified hydrophobic surface exposure in CDR regions (SAP_pos_CDR) as the dominant predictor, followed by heavy-chain variable domain SAP; charge-based descriptors contributed minimally. These CDR loops are key determinants of both binding specificity and developability risk [24, 25], suggesting that VIBE1 captures non-specific interaction propensity driven primarily by hydrophobic patch exposure. The residual variance unexplained by static sequence features suggests VIBE1 additionally encodes information from conformational dynamics and collective interfacial behaviour [29, 30].

## Discussion

Developability methods have historically been motivated by one of two complementary strategies. The first is mechanistic, encompassing both methods that simulate a stress encountered during the lifecycle of a therapeutic molecule (as in agitation-stress tests, where mechanical perturbation directly probes aggregation susceptibility) and methods that probe underlying molecular attributes likely to become liabilities under such stresses (HIC for hydrophobicity, SINS for self-interaction). Interfacial adsorption and mechanical disruption during manufacturing, fill-finish, and IV administration are well-documented drivers of protein aggregation and particle formation [39], and extensional flow during syringe and pump operations subjects molecules to elongational forces that can unfold and aggregate even thermodynamically stable proteins [40]. The second strategy is empirical: exposing molecules to a controlled environment chosen for discriminating power rather than direct in-vivo relevance, and using a data-driven approach to identify which responses correlate with developmental outcomes. The BVP polyreactivity assay measures binding to baculovirus particles, a complex biological surface that is not a direct mimic of any single in vivo liability but may present interaction modes relevant to clearance behaviour; this approach has proven predictive of rapid clearance and poor pharmacokinetics [2]. Jain et al. demonstrated that combining readouts from both types of method into a multiparameter framework substantially improved discrimination between molecules that progressed clinically and those that did not [3]. The LSI measurement draws on both traditions. Each droplet event exposes molecules to interface creation and annihilation, elongational stress during pinch-off [40, 41], and abrupt changes in ionic and excipient environment, stresses that occur, among other contexts, during IV bag preparation and infusion, where surfactant dilution and air-liquid interface exposure are recognised risks for aggregation [39]. At the same time, the high-dimensional behavioural fingerprint it generates is not designed to isolate any single mechanism, but to capture composite responses whose predictive value, like that of BVP, is established by correlation with outcomes.

We have introduced the VIBE method, executed via the LSI sensing architecture, as a new modality for early-stage biophysical profiling of therapeutic antibodies. The primary descriptor, VIBE1, integrates information across multiple biophysical dimensions into a single behavioural readout derived from interfacial wave dynamics. By operating at less than 10 *µ*g of material per measurement, this approach breaks the three-way trade-off between data quality, material efficiency, and measurement speed that has historically confined behavioural assessment to late-stage development.

### Relation to existing methods

A key finding is that VIBE1 does not recapitulate any single conventional method. Across eleven established methods spanning hydrophobicity, polyreactivity, self-interaction, thermal stability, and aggregation propensity, VIBE1 showed individually modest correlations (*ρ* ≈ 0.2–0.3), yet positioned centrally between method clusters in hierarchical analysis [21, 16]. This pattern is consistent with an integrative behavioural signal rather than a proxy for any known single property. Accordingly, when ranked by discriminative power for clinical outcome on the Jain et al. overlap cohort, VIBE1 placed within the modest discrimination range achieved by several widely used methods, despite requiring orders of magnitude less material and no method-specific reagents.

### Progressive filtering across development stages

The fraction of high-VIBE1 antibodies declined monotonically from Phase I (20.7%) to approved drugs (9.2%), mirroring the pattern reported by Jain et al. [3, 21] for biophysical outliers. This implies that the risk captured by VIBE1 manifests throughout the development lifecycle: a measurement available at hit identification, from micrograms of material, captures risk that currently becomes visible only through cumulative pipeline attrition.

### High-precision triage at hit identification

The operating characteristic of VIBE1, high precision and moderate recall, matches the asymmetric cost structure of early-stage triage, where incorrectly discarding a viable candidate is costlier than missing a problematic one that downstream methods may catch. VIBE1 achieved no false positives among the 6 antibodies it flagged in the industrial cohort (precision 100%, 95% CI: 61–100%) and a precision of 85% on the 41 it flagged in the clinical cohort (95% CI: 72–93%), with recall of approximately 33% and 21% respectively. This positions it as a reliable first-pass filter that is additive to existing workflows: it does not catch every problematic molecule, but those it flags can be deprioritised with high confidence.

### Sensing principle

The LSI architecture extends the perturbation-based measurement tradition established by Eigen [19] to interfacial dynamics: kinetic differences between molecules are resolved by the system’s dynamic response to controlled excitation rather than by equilibrium partitioning. The substrate is operated near a thermodynamic transition, a regime in which dynamical signals are predicted to be amplified and structured [13, 33, 14, 32]; the computational framing of hydrodynamic reservoirs motivates the broader sensing architecture [11, 12].

### Practical utility

The VIBE method operates at 0.1 mg/mL with approximately 80 *µ*L sample volumes, corresponding to less than 10 *µ*g of material per measurement. This makes it compatible with hit identification and early lead optimisation, where material is limiting and the candidate pool is largest. By generating integrated behavioural data at this stage, months before conventional developability characterisation becomes feasible, the approach repositions molecular behaviour from a late-stage validation checkpoint to a discovery-phase design variable.

### Robustness against overfitting

The descriptor selection (ANOVA *F*-statistic on biophysical-flag classes) and threshold setting (two-component GMM on replicate-averaged VIBE1 values) were both performed without exposing the clinical-outcome labels to the analysis pipeline. The top descriptor coincided across Dataset 1 and Dataset 2, and a tightly correlated family of high-*F* features (SI Appendix, Figure S1) indicates the signal is not carried by a single fragile descriptor. Larger and prospectively-collected validation cohorts remain desirable.

### Limitations

The sensing liquid composition and signal-processing parameters are proprietary, constraining independent reproduction at the hardware level; descriptor-level outputs are publicly available. VIBE1 does not provide a mechanistic diagnosis of specific failure modes, and validation is currently limited to monoclonal antibodies.

## Materials and Methods

### Antibody samples

Dataset 1 comprised 71 antibodies provided by Boehringer Ingelheim (1 mg/mL in 10 mM histidine, pH 6.0, 20 mM NaCl) and diluted to 0.1 mg/mL in PBS for VIBE analysis. Dataset 2 comprised 235 clinical-stage therapeutic antibodies sourced from Sino Biological (HEK293; supplied in PBS, pH 7.4) and matched to the corresponding clinical sequences and reported constant-region isotypes in Thera-SAbDab [20] (SI Appendix, Table S1); Dataset 2 therefore includes variable isotypes as used clinically. For concordance analyses against the eleven-method panel of Jain *et al*. [21], the overlap set comprised 135 antibodies of the IgG1 isotype: 82 that were already IgG1 in Dataset 2, plus 53 non-IgG1 antibodies re-expressed in IgG1 format to ensure like-for-like comparison. VIBE1 values for the 53 re-expressions were taken from their IgG1 measurements; values for the 82 native-IgG1 antibodies were taken from the original Dataset 2 runs.

### LSI measurement

Samples (0.1 mg/mL, 80 *µ*L injection) were deposited droplet-by-droplet onto a proprietary near-thermodynamic-transition sensing liquid [11]. Interfacial waves were recorded via ellipsometry at 100 kS/s for ~400 s per sample, yielding up to 200 million data points per antibody. Samples were incubated at 4°C for 48 hours prior to testing to mitigate time-dependent adsorption effects and tested in randomised-order batches of 20, run in triplicate, at 22°C *±* 2°C. Between measurements, the trough was cleaned with 10% 2-propanol followed by re-equilibration with deionised water.

### VIBE1 descriptor

VIBE1 is defined by a two-pass hierarchical moment analysis over the interaction window (droplets 20–40) and signal window (104–140 ms). First pass: 4th central moment across droplets at each time point, root-standardised (Eqs. S1–S2). Second pass: mean across time points (Eq. S4). Selected from 320 candidates by ANOVA *F*-statistic; independently top-ranked in both datasets. Full parameter-search details in SI Appendix.

### Classification and statistical analysis

A two-component GMM defined the threshold *τ*_2_ = 0.041 on Dataset 2, transferred to Dataset 1 via min–max rescaling against sirukumab/trastuzumab controls (Eqs. S5–S6; *τ*_1_ = 0.036). Twosided 95% confidence intervals for precision, recall, failure rates, and other binomial proportions were computed using Wilson score intervals [42]. Confidence intervals for the difference between two proportions (e.g. flagged vs. baseline failure rate) were computed using the Newcombe method based on Wilson intervals. Uncertainty for ROC-AUC in the structural analysis was assessed by non-parametric bootstrap (1,000 iterations) on out-of-fold predicted probabilities. Correlations: Spearman’s *ρ*. Structural analysis: Random Forest on DeepSP descriptors with SHAP. Full details in SI Appendix.

## Acknowledgments

We thank Brian Fiske and Nimish Gera of Mythic Therapeutics for their early conviction in the potential of our technology and for their thoughtful input during its formative stages. Their engagement helped us better understand how emerging biophysical platforms must align with practical decision-making in therapeutic development. We are also especially grateful to Pete Tessier for his thoughtful review of the manuscript.

