## Supplemental Information for "Behavioural Profiling of Therapeutic Antibodies via Non-equilibrium Interfacial Wave Dynamics"

### Supporting Information for:
### Behavioural Profiling of Therapeutic Antibodies via Non-equilibrium Interfacial Wave Dynamics

Alec Thomas[1], Alexander St John[1], Maureen Crames[2], Piergiorgio Caramazza[1], Alireza Meghdadi[1], Joe Bailey[1], Rodolfo Hermans[1], Sejeong Lee[1], Elizabeth Lam[1], Vivek Krishnan[1], Alexandra Amoruso[1], Sam Aoudjane[1], Michael S. Marlow[2], Carolina Zirn[1], Pratik Kotkar[1], Akshay Mishra[1], Anshika Srivastava[1], and Shamit Shrivastava*[1]

[1]Apoha Limited, 135 Salusbury Road, NW6 6RJ, London, UK
[2]Biotherapeutics Discovery, Boehringer Ingelheim Pharmaceutical Inc, 900 Ridgebury Road, Ridgefield, CT 06877, USA

#### Materials and Methods

##### Antibody samples

A total of 306 unique antibodies were characterised across two primary datasets (Table S1), with an additional 53 re-expressed as described below. Of the 135 clinical-stage antibodies in Dataset 2 that overlapped with the study by Jain et al. [3], 82 were already of the IgG1 isotype. An additional 53 antibodies of non-IgG1 isotype were re-expressed in IgG1 format to enable concordance comparison with the Jain et al. method panel, which used IgG1 throughout. Together, these 135 antibodies (82 + 53) constitute the concordance subset. The 53 re-expressions were formulated identically to Dataset 2 (PBS, pH 7.4) and measured under the same VIBE conditions; they were excluded from GMM fitting and clinical enrichment analyses.

**Dataset 1** comprised 71 antibody samples provided by Boehringer Ingelheim at approximately 1 mg/mL in 10 mM histidine (pH 6.0) and 20 mM NaCl. These were accompanied by measurements from five standard developability methods (purity after Protein A capture by SEC (% purity), Fab $T_m$, aHIC, SINS, NSB) [1, 2, 3]. For VIBE analysis, stock samples were diluted 10-fold in PBS to 0.1 mg/mL. Like Dataset 2, Dataset 1 was dominated by IgG1-format antibodies, providing a consistent isotype basis for direct cross-dataset comparison.

**Dataset 2** comprised 235 recombinantly expressed antibodies produced by Sino Biological (Beijing, China) using HEK293 cells, formulated in PBS (pH 7.4). Stock concentrations ranged from 0.3 to 9 mg/mL; samples were stored at −80°C until use. For VIBE analysis, stock samples were diluted to a final concentration of 0.1 mg/mL in PBS. Clinical status and phase-progression history were sourced from Thera-SAbDab [4]. Full sequence information is provided below.

#### Sample preparation and deposition

Antibody samples were thawed overnight at 4°C and diluted to 0.1 mg/mL in PBS (pH 7.4). For each analysis, 300 $\mu$L of antibody solution was aliquoted into glass HPLC vials and maintained at 4°C in the autosampler. Sample deposition was driven by an HPLC system (without chromatographic column) under the following conditions: 80 $\mu$L injection volume, 100 $\mu$L/min flow rate of deionised water, isocratic elution, 7-minute run time, with UV absorbance monitored at 280 nm for quality verification.

#### LSI measurement

Within the LSI module, antibody samples were deposited droplet-by-droplet onto an engineered liquid trough containing a proprietary sensing liquid, a surface-active colloidal medium maintained near a thermodynamic phase transition [5]. Each droplet deposition initiates interfacial interactions, generating an interfacial wave response that combines contributions from capillary and gravity modes; this response is recorded via an ellipsometric optical setup with 16-bit resolution at 100 kS/s sampling rate for approximately 400 s per sample, yielding up to 200 million data points per antibody. After each measurement, the trough undergoes automated cleaning and refilling.

The system was maintained at 22°C ± 2°C. Samples were tested in randomised-order batches of 20 and run in triplicate. Between measurements, a cleaning protocol of 10% HPLC-grade 2-propanol (1.5 min) followed by re-equilibration with mobile phase (purified deionised water) was employed to prevent cross-contamination. Samples were incubated in their testing vials at 4°C for 48 hours prior to testing to mitigate time-dependent adsorption effects.

#### VIBE descriptor framework and parameter search

Let $\mathbf{X}$ denote the waveform matrix formed by selecting $D = 21$ droplets from the interaction window (droplets 20–40; rows, indexed by $i = 1, \ldots, D$) and $T$ time points within the chosen signal window (columns, indexed by $j = 1, \ldots, T$). Each entry $x_{ij}$ represents the signal amplitude at time point $t_j$ for the $i$-th droplet in the selected optical channel.

In the first pass, the $n$-th central moment is computed column-wise:

$$\hat{\mu}_{n,j} = \frac{1}{D} \sum_{i=1}^{D} (x_{ij} - \bar{x}_j)^n \tag{S1}$$

where $\bar{x}_j = \frac{1}{D} \sum_{i=1}^{D} x_{ij}$ and $n \in \{1, 2, 3, 4\}$. The $n$-th root standardisation is then applied:

$$\mu^*_{n,j} = \operatorname{sgn}(\hat{\mu}_{n,j}) \cdot |\hat{\mu}_{n,j}|^{1/n} \tag{S2}$$

In the second pass, the $m$-th central moment across time points yields the raw descriptor:

$$\mathrm{VIBE}_{n,m} = \frac{1}{T} \sum_{j=1}^{T} \left( \mu^*_{n,j} - \bar{\mu}^*_n \right)^m \tag{S3}$$

where $\bar{\mu}^*_n = \frac{1}{T} \sum_{j=1}^{T} \mu^*_{n,j}$ and $m \in \{1, 2, 3, 4\}$. For $m \geq 2$, Eq. S3 computes the $m$-th central moment across time points. For $m = 1$, the first central moment is identically zero, so the expression reduces by convention to the arithmetic mean: $\mathrm{VIBE}_{n,1} = \bar{\mu}_n^*$.

The systematic scan covered four optical channels ($S_Y$, $S_{\mathrm{Sum}}$, $P$, $S_X$), five signal windows (98–104 ms, 90–140 ms, 90–98 ms, 104–140 ms, 90–104 ms), and all 16 moment-pair combinations ($n, m \in \{1, 2, 3, 4\}$), yielding 320 candidate features. Each was evaluated by ANOVA $F$-statistic across antibody identities. Features above $F \approx 20$ exhibited absolute Spearman correlations exceeding 0.8 with the top-ranked feature (Figure S1). The top-ranked feature, VIBE1, is:

$$\mathrm{VIBE1}_{\mathrm{raw}} = \frac{1}{T} \sum_{j=1}^{T} \mu_{4,j}^* \tag{S4}$$

computed on the $S_Y$ channel with signal window 104–140 ms. The same feature was independently top-ranked in Dataset 2.

#### Classification threshold and cross-dataset transfer

Because Dataset 2 was acquired first and comprises a larger, more diverse clinical cohort ($n = 235$), the classification threshold was defined on this dataset. A two-component Gaussian mixture model (GMM) was fitted to replicate-averaged VIBE1 values from the 235-antibody clinical cohort. The 53 IgG1 re-expressions described above were excluded from GMM fitting and clinical enrichment analyses. The GMM identified a dominant component and a minority component; the threshold $\tau_2 = 0.041$ was defined as the intersection of the two weighted densities (Figure S2C-D).

To transfer the threshold to Dataset 1, which was acquired under different buffer conditions (histidine vs. PBS), $\tau_2$ was expressed as a fractional position within the sirukumab–trastuzumab control interval in Dataset 2:

$$\hat{\tau} = \frac{\tau_2 - \bar{v}_{\mathrm{tras}}^{(2)}}{\bar{v}_{\mathrm{siru}}^{(2)} - \bar{v}_{\mathrm{tras}}^{(2)}} \tag{S5}$$

and projected onto the Dataset 1 control interval:

$$\tau_1 = \hat{\tau} \cdot (\bar{v}_{\mathrm{siru}}^{(1)} - \bar{v}_{\mathrm{tras}}^{(1)}) + \bar{v}_{\mathrm{tras}}^{(1)} \tag{S6}$$

yielding $\tau_1 = 0.036$. A *de novo* GMM fitted directly to Dataset 1 produced an almost identical threshold of 0.037 (Figure S2A-B), confirming cross-dataset consistency.

#### Confidence intervals

Confidence intervals for binomial proportions (precision, recall, failure rates, flagged fractions) were computed using Wilson score intervals [8]. Confidence intervals for the difference of two independent proportions (e.g. flagged failure rate versus baseline) were computed using the Newcombe method with Wilson endpoints. For the DeepSP structural classifier (see below), 95% confidence intervals for ROC-AUC were obtained by non-parametric bootstrap resampling ($n = 1{,}000$ iterations) of out-of-fold predicted probabilities, with the 2.5th and 97.5th percentiles defining the interval.

#### Clinical enrichment metrics

Clinical failure rate was defined as the fraction of non-approved antibodies within the flagged subset; baseline failure rate was defined over the full cohort. Enrichment: ratio of flagged to baseline failure rate. Relative success-rate uplift: $(p_{\text{kept}} - p_{\text{base}})/p_{\text{base}}$. Confidence intervals for failure rates and flagged fractions were computed using Wilson score intervals and the Newcombe method, as described in the Confidence intervals section above.

#### Correlation analyses

For Dataset 2, the eleven-method concordance heatmap was computed on the IgG1 concordance subset ($n = 135$; 82 native IgG1 plus 53 re-expressed), which share variable domains with a subset of the clinical cohort but were measured as a matched IgG1 format specifically for this comparison. Pairwise Spearman rank correlations were computed between per-antibody mean VIBE1 and conventional method measurements. Statistical significance was assessed at $\alpha = 0.05$. Trend lines were fitted by ordinary least squares for visualisation only.

#### Structural descriptors

SAP and SCM descriptors were computed from antibody variable domain sequences using DeepSP [6]. A Random Forest classifier (scikit-learn, $n\_estimators = 100$) was trained to distinguish high- from low-VIBE1 antibodies using all descriptors as inputs. Performance was evaluated by stratified 5-fold cross-validation (`shuffle=True`, `random_state` = 42), with out-of-fold predicted probabilities concatenated across folds. Bootstrap confidence intervals ($n = 1{,}000$) were computed on these out-of-fold probabilities. SHAP values [7] were computed using a single Random Forest fitted to the full dataset.

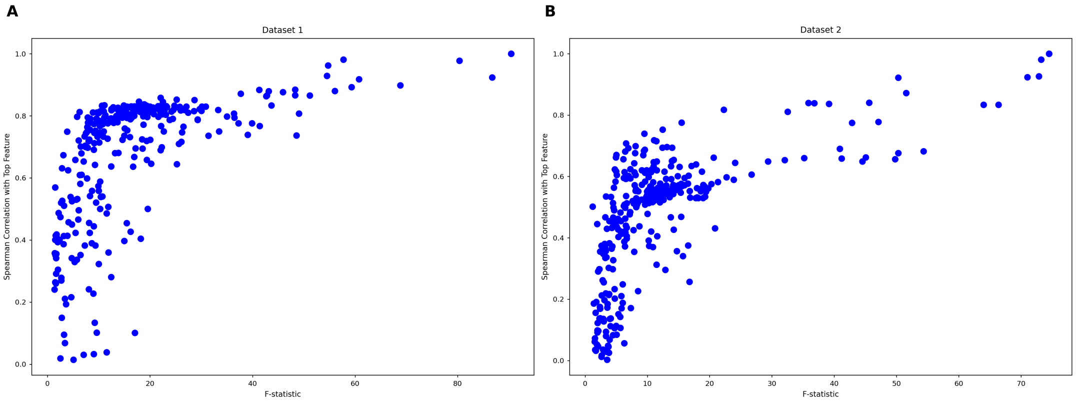


**Figure S1. High-$F$ features form a correlated cluster in both datasets.** Spearman correlation with the top-ranked feature plotted against ANOVA $F$-statistic for all 320 candidate features. ($A$) Dataset 1. ($B$) Dataset 2. In both datasets, features with increasing discriminatory power (higher $F$-statistic) converge towards strong positive correlation with the top-ranked descriptor, VIBE1, forming a dense upper envelope. This pattern indicates that the highest-ranked features constitute a coherent, tightly correlated cluster capturing the same dominant behavioural signal, rather than representing isolated or unstable optima in the feature search space.

**Table S1. Dataset 2 antibody panel.** Clinical-stage therapeutic antibodies ($n = 235$) with matched Thera-SAbDab identifiers, constant-region isotype, highest clinical phase reached, and current development status. Antibodies are listed alphabetically by INN.

| INN | Thera-SAbDab ID | Isotype | Highest Phase | Status |
|---|---|---|---|---|
| *[DATA TO BE INSERTED]* | | | | |

**Abbreviations:** INN, International Nonproprietary Name. Highest Phase: I, II, III, or Approved. Status: Active, Discontinued, or No Further Development. Isotype as reported in Thera-SAbDab [4]. Clinical status current as of database access date.

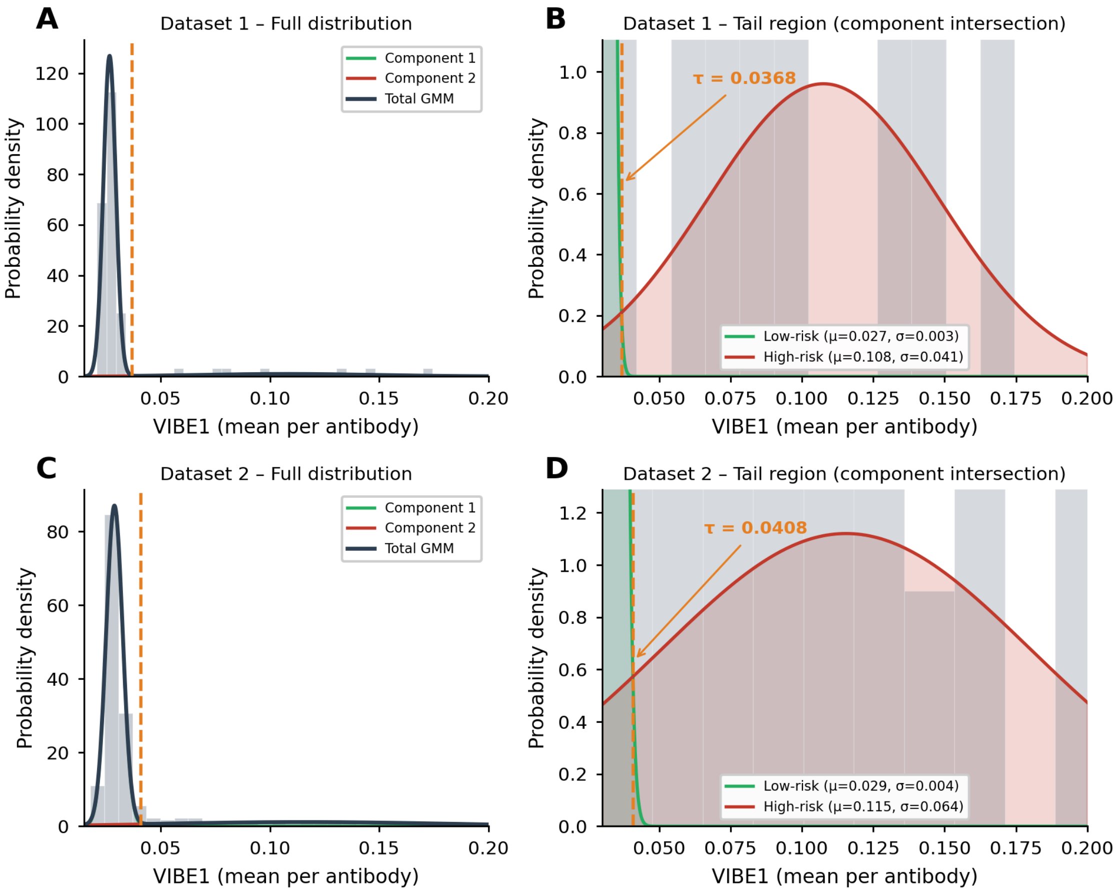


**Figure S2. Gaussian mixture model (GMM) threshold estimation for VIBE1.** (*A*) Dataset 1: two-component GMM fitted to replicate-averaged VIBE1 values (all samples including controls), shown for comparison to the transferred threshold. Green curve: low-VIBE1 component; red curve: high-VIBE1 component; dark blue curve: total mixture density. Grey bars: histogram of observed VIBE1 values. (*B*) Dataset 1: zoomed view of the tail region and component intersection, yielding a consistent threshold ($\tau = 0.0368$). Low-VIBE1 component ($\mu = 0.027$, $\sigma = 0.003$); high-VIBE1 component ($\mu = 0.108$, $\sigma = 0.041$). (*C*) Dataset 2: two-component GMM fitted to replicate-averaged VIBE1 values across 235 antibodies. Dashed orange line: classification threshold ($\tau = 0.0408$). (*D*) Dataset 2: zoomed view of the tail region showing the component intersection used to define $\tau$. Low-VIBE1 component ($\mu = 0.029$, $\sigma = 0.004$); high-VIBE1 component ($\mu = 0.115$, $\sigma = 0.064$).

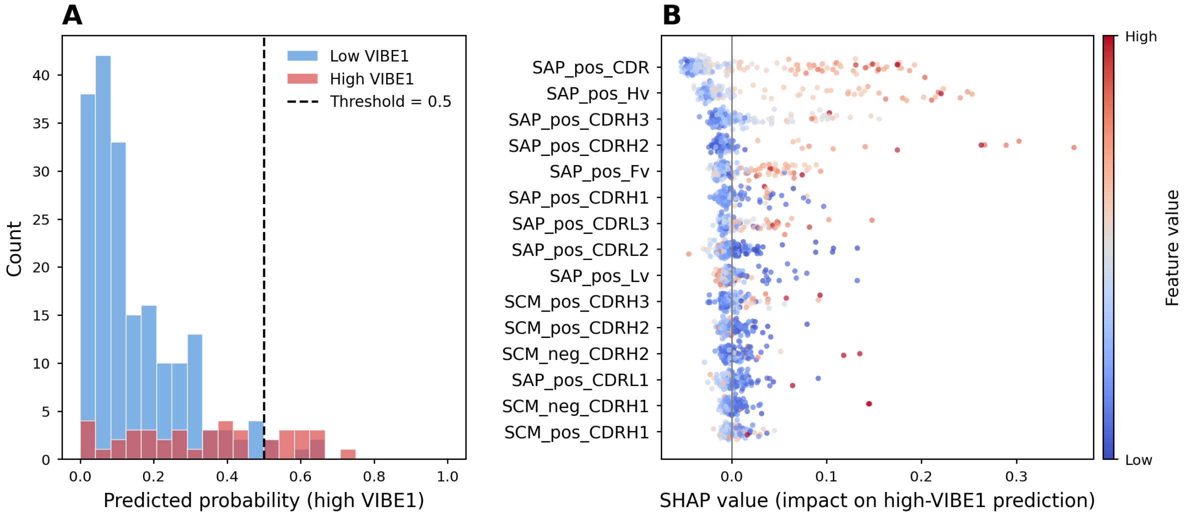


**Figure S3. DeepSP structural descriptors predict high-VIBE1 antibodies. (A)** Predicted probability distributions from the Random Forest classifier for low-VIBE1 (blue) and high-VIBE1 (red) antibodies. Dashed line: decision threshold of 0.5. ROC-AUC = 0.766 (95% CI: 0.665–0.851). **(B)** SHAP summary plot. Features ranked by mean absolute SHAP value; points coloured by feature value (red = high, blue = low). Hydrophobic surface exposure in CDR regions (SAP__pos__CDR) is the dominant predictor; charge-based descriptors (SCM) contribute minimally.
